# Links between central CB1-receptor availability and peripheral endocannabinoids in patients with first episode psychosis

**DOI:** 10.1101/664086

**Authors:** Alex M. Dickens, Faith Borgan, Heikki Laurikainen, Santosh Lamichhane, Tiago Marques, Tuukka Rönkkö, Mattia Veronese, Tuomas Lindeman, METSY Investigators, Tuulia Hyötyläinen, Oliver Howes, Jarmo Hietala, Matej Orešič

## Abstract

There is an established, albeit poorly-understood link between psychosis and metabolic abnormalities such as altered glucose metabolism and dyslipidemia, which often precede the initiation of antipsychotic treatment. It is known that obesity-associated metabolic disorders are promoted by peripheral activation of the endocannabinoid system (ECS). Our recent data suggest that ECS dysregulation may also play a role in psychosis. With the aim of characterizing the involvement of the central and peripheral ECSs and their mutual associations, here we performed a combined neuroimaging and metabolomic study in patients with first-episode psychosis (FEP) and healthy controls (HC). Regional brain cannabinoid receptor type 1 (CB1R) availability was quantified in two, independent samples of patients with FEP (n=20 and n=8) and HC (n=20 and n=10), by applying 3D positron emission tomography (PET), using two radiotracers, [^11^C]MePPEP and [^18^F]FMPEP-d2. Ten endogenous endocannabinoids or related metabolites were quantified in serum, drawn from these individuals during the same imaging session. Circulating levels of arachidonic acid and oleyl ethanolamide were reduced in FEP individuals, but not in those who were predominantly medication-free. In HC, there was an inverse association between levels of circulating arachidonoyl glycerol, anandamide, oleyl ethanolamide and palmitoyl ethanolamide, and CB1R availability in the posterior cingulate cortex. This phenomenon was, however, not observed in FEP patients. Our data thus provide evidence of cross-talk and dysregulation between peripheral endocannabinoids and central CB1R availability in FEP.

## Introduction

Psychotic disorders are associated with a reduced life expectancy of 15-20 years^1^, which is in part due to the high prevalence of cardiovascular disease, type 2 diabetes (T2DM) and metabolic syndrome^2, 3^. Unhealthy lifestyles and pharmacological side effects have been suggested to be a major cause of these mortality rates^4^. However, abnormal glucose homeostasis, hyperinsulinemia, dyslipidemia and accumulation of visceral fat are already evident in drug-naïve first episode psychosis (FEP) patients, independent of obesity^5, 6^.

The mechanisms underlying the development of metabolic co-morbidities in psychosis are poorly understood. There is evidence that anti-psychotic medication, particularly clozapine, leads to rapid weight gain^7^, while co-treatment with metformin can help reduce this^8^. A recent study in FEP patients identified changes in specific circulating lipids, which are known to be associated with non-alcoholic fatty liver disease (NAFLD) and T2DM risk, prior to weight gain^9^.

There is emerging evidence suggesting that the endocannabinoid system (ECS) might be dysregulated in various psychiatric disorders, including schizophrenia^10–14^. Furthermore, the ECS has been shown to play a role in metabolic disorders^15–17^. Taken together, these findings suggest that the dysregulation of the ECS could play a role in the development or progression of metabolic co-morbidities observed in psychosis.

The ECS is composed of (1) cannabinoid receptors type 1 (CB1R) and type 2 (CB2R), (2) lipid-derived endocannabinoid ligands with affinity to these or other receptors, and (3) enzymes involved in the synthesis and degradation of these ligands^18, 19^. The CB1Rs are found in the central nervous system (CNS) as well as in the periphery, throughout the gastrointestinal tract, liver, adipose tissue and adrenal glands^20, 21^. In the CNS, the CB1R plays an important role in cognition^22^ and has been implicated in mood and anxiety disorders^23, 24^. Recent evidence from both our group and others suggests that central CB1R availability is altered psychosis^25–27^. We recently showed that CB1R availability is altered in patients with psychosis, where greater reductions in CB1R levels are associated with greater symptom severity and poorer cognitive functioning^27^. There is also evidence that endogenous cannabinoid ligands which act as CB1R agonists are elevated in the cerebrospinal fluid of medication-naïve psychotic patients^11, 14, 28, 29^.

Metabolomics approaches have identified systemic metabolic changes in the context of psychosis^9, 30, 31^ and neurodegenerative disorders^32–34^, suggesting that a strong link exists between the circulating metabolome and diseases of the central nervous system. Several studies have demonstrated that psychotic patients have dysregulated metabolic profiles, including dyslipidemia and dysglycemia^35, 36^. Intriguingly, an imbalance of peripheral endocannabinoids has also been associated with various metabolic disorders, including diabetes and obesity^37, 38^. Furthermore, changes in blood endocannabinoid levels are associated with improvement in symptoms of psychosis^39^, which could imply that endocannabinoid signaling may not only be dysregulated in the central nervous system of psychotic patients, but also in the periphery^12^. There is an association between cannabis use and disturbances of the central ECS, providing further evidence for potential cross-talk between the endogenous peripheral and central ECSs^40, 41^. However, to our knowledge, there are no data investigating the link between peripheral endocannabinoid levels and brain CB1R availability in healthy individuals or in psychosis.

Therefore, we aimed to investigate the association between peripheral endocannabinoids and brain CB1R availability. To this end, we quantified serum endocannabinoids using liquid-chromatography triple-quadrupole mass spectrometry (LC-QqQMS) and brain CB1R availability in a case-control setting, which included FEP patients and matched healthy controls.

## Methods

### Ethics statement

Ethical approvals were obtained from respective study sites in Finland (ETMK 98/180/2013) and England (14/LO/1289). Subjects’ capacities for consent were assessed and informed, written consent was obtained from all volunteers.

### Study design and participants

The study setting was described in detail previously^27^. CB1R availability was investigated at two PET centers using independent samples; Turku (n = 11, HC; n = 8, FEP) and London (n = 23, HC; n = 20, FEP). However, some subjects were excluded due to increases in AG levels, caused by sample handling resulting in the following subjects being included in the analysis: Turku (n = 10, HC; n = 8, FEP) and London (n = 15, HC; n = 11, FEP) (Table 1). Given putative sex differences in CB1R availability^42^, we only investigated males to remove sex as a source of variability. Corresponding studies in females are currently ongoing.

**Table 1.** Demographic characteristics of the study populations. *p<0.05

|  | Turku |  | KCL |  |
| --- | --- | --- | --- | --- |
|  | FEP<br>n = 8<br>mean<br>(SD) | HC<br>n = 10<br>mean (SD) | FEP<br>n = 15<br>mean (SD) | HC<br>n = 11<br>mean (SD) |
| Age (years) | 26.4 (3.6) | 27.18 (5.9) | 27.1 (5.1) | 26.6 (6.9) |
| BMI (kg/m<sup>2</sup>) | 28.2 (6.9) | 25.3 (3.7) | 25.9 (5.2) | 26.3 (4.1) |
| Years of education | 13.3 (1.3) | 15.7 (3.2) | n/a | n/a |
| Illness duration<br>(months) | 5.4 (6.8) | n/a | 27.1 (10.1) | n/a |
| BPRS total score sum | 65.3 (17.6)* | 30.6 (2.4)* | n/a | n/a |
| BPRS positive score sum | 20.0 (7.3)* | 9.2 (0.6)* | n/a | n/a |
| PANSS positive | n/a | n/a | 24.4 (5.4) | n/a |
| PANSS negative | n/a | n/a | 42.5 (9.0) | n/a |
| PANSS general | n/a | n/a | 42.53 (9.0) | n/a |
| PANSS total | n/a | n/a | 90.7 (16.6) | n/a |

FEP patients met the following inclusion criteria: (i) DSM-IV diagnosis of a psychotic disorder, determined by the SCID-I/P; (ii) illness duration of less than 3 years; and (iii) male sex. In Turku, FEP volunteers were already taking antipsychotic medication prior to the start of the study and had diagnoses of affective or non-affective psychosis. In London, FEP volunteers were, with the exception of two individuals, medication-free from all pharmacological treatments for at least 6 months and had diagnoses of schizophrenia or schizoaffective disorder (DSM-IV criteria). Healthy volunteers had no current/lifetime history of any Axis-I disorder as determined by the SCID-I/P and were matched for age (age +/-3 years) and sex (male). Exclusion criteria for all volunteers were: (i) current/lifetime history of substance abuse/dependence; (ii) substance use within the last month; (iii) positive screen on toxicology tests for cannabis and other substances.

### CB1R availability PET imaging

The full methods for PET CB1R imaging were described in detail previously^27^. However, a concise description for both studies, Turku and London, is provided here.

#### Turku study

[^18^F]FMPEP-d2 was synthesized as described previously^43^, with slight modifications. The radiochemical purity was greater than 95% and the molar radioactivity was greater than 500 GBq/μmol at the end of synthesis resulting in an injected tracer mass below 200 ng.

Positron emission scans were done in three-dimensional list-mode, in increasing frame duration for a total scan range of 0-120 min, with a brain-dedicated high-resolution research tomograph (ECAT HRRT, Siemens Medical Solutions, Kemnath, Germany), after a bolus injection of [^18^F]FMPEP-d2 (201 ± 11.1 MBq). Attenuation correction of PET was applied using a ^137^Cs point source. Arterial plasma activity was measured continuously for the first 3.5 minutes, and discretely thereafter at 4.5, 7.5, 11,15, 20, 25, 30, 35, 40, 45, 50 and 60 min. Arterial input was corrected for tracer metabolite activity measured using thin layer chromatography and digital autoradiography. The resulting metabolite free arterial input was corrected for temporal delay between blood and PET tissue measurements. Motion correction between PET frames was done by realigning the PET frames to a single reference frame with the most uptake on average. Whole-brain T1-weighted MRI images were acquired by a Philips 3T Ingenuity PET/MR hybrid scanner. The individual T1-weighted images were co-registered to the sum of the realigned PET frames. Inverse normalization parameters obtained from normalizing the co-registered T1-weighted images to MNI space were used to fit the Hammersmith anatomical atlas to the PET frames (Hammers et. al 2003). Data pre-processing was performed using Statistical Parametric Mapping 12 (http://www.fil.ion.ucl.ac.uk/spm) and MATLAB R2014b (Mathworks Inc., Natick, MA).

#### London study

Continuous, 90-minute PET scans were acquired on a PET/CT (Hi-Rez Biograph 6 CT44931; Siemens Medical Solutions, Kemnath, Germany) in three-dimensional mode, after bolus injection of 314 ± 34.4 MBq of [^11^C]MePPEP, as previously described^44–46^. CT scans were acquired prior to each PET scan for correction of attenuation and scatter. Continuous arterial blood sampling took place for the first 15 minutes of the scan which was followed by discrete blood sampling at 2, 5, 10, 15, 20, 25, 35, 40, 50, 60, 70, 80 and 90 minutes after radioligand injection. Images were reconstructed with filtered back-projection including corrections for attenuation and scatter. There were no significant group differences in injected mass, injected activity, or specific activity. To aid anatomical localization, each volunteer received a high-resolution structural 3D T1-weighted magnetic resonance scan acquired on a General Electric MR750 3T scanner (MR750; GE Healthcare, Chicago, IL).

Data pre-processing was performed using a combination of Statistical Parametric Mapping 12 (http://www.fil.ion.ucl.ac.uk/spm) and FSL (http://www.fsl.fmrib.ox.ac.uk/fsl) functions, as implemented in MIAKAT (http://miakat.org). Motion correction was applied to non-attenuation-corrected images^47^. Non-attenuated corrected frames were realigned to a single “reference” frame (corresponding to that with the highest number of counts) by employing a mutual information algorithm. The transformation parameters were then applied to the corresponding attenuated-corrected dynamic images, creating a movement-corrected dynamic image which was used for subsequent analysis. Realigned frames were then summated to create single-subject motion-corrected maps which were then used for MRI and PET co-registration, prior to PET data quantification. T1-weighted structural images were co-registered to the PET image using rigid body transformation. Normalization parameters were obtained by warping the co-registered structural MRI to MNI space (International Consortium for Brain Mapping ICBM/MNI) using bias-corrected segmentation. The inverse of these parameters was used to fit a neuroanatomical atlas to each individual PET scan using the Hammersmith atlas^48^. Whole blood time-activity curves (TACs) were fitted using a multi-exponential function as derived by Feng’s model^49^. For each scan, a time delay was fitted and applied to the input functions (both parent and whole blood TACs) to account for any temporal delay between blood sample measurement and target tissue data. Regions of interest were harmonized with the Turku PET dataset.

In both Turku and London studies, CB1R availability was indexed using the volume of distribution (VT ml/cm^3^) of the respective PET tracer, as calculated using the Logan graphical method with a metabolite-corrected arterial plasma input function. The resulting model was validated as described previously^27, 50^. The temporal lobe, frontal lobe, anterior cingulate cortex, posterior cingulate cortex, striatum, parietal lobe, occipital lobe, putamen, nucleus caudatus, insula, amygdala, and whole brain were chosen as regions of interest for PET. Hippocampal, amygdala, superior temporal gyral, superior frontal, middle frontal, inferior frontal and orbitofrontal subregions were included to explore regional differences within relevant regions of interest (ROI). Due to the use of different PET tracers used in both sites, the CB1R availability data was scaled to unit variance and zero mean within each study, when analyzed together.

### Analysis of serum endocannabinoids

The following peripheral endocannabinoids and related structures were measured using a targeted LC-QqQMS) method: THC-COOH, arachidonoyl ethanolamide (AEA), N-arachidonoyl dopamine (NADA), 2-arachindonoyl glycerol (2-AG), 1-arachidonoyl glycerol (1-AG), oleyl ethanolamide (OEA), palmitoyl ethanolamide (PEA), 2-arachidonoyl glycerol ether (2-AGe), arachidonic acid (AA) and stearoyl ethanolamide (SEA). The analytical method was developed for serum samples and all solvents were liquid chromatography (LC) grade (Sigma-Aldrich Inc., St. Louis, MO). Initially, the internal standards (10 µL, 100 ng/mL, of THC-COOH-d9, 2-AG-d5, NADA-d8, and AEA-d8 in EtOH) were added to 200 µL of serum in silylated 2 mL glass chromatography vials. The proteins were then precipitated using acetonitrile (400 µL) containing 0.1% v/v formic acid (LC-grade, Sigma-Aldrich). This was done in metal and glass syringes to ensure no contamination from plastic. The samples were then briefly vortexed and placed at −20 °C for 30 minutes. The samples were then centrifuged (10,000 g, 4 °C) for 10 minutes. The supernatant was then removed and placed in fresh silylated vials and stored for no more than 24 hours before solid phase extraction (SPE) of the endocannabinoids.

The solid-phase extraction (SPE, reverse phase HLB, 30 mg sorbent) was performed in 96-well plates (Waters Inc., Milford, MA) and performed on ice to reduce plastic contamination. Initially, 1 mL of ultrapure H_2_O was added to the top of each well. The supernatant was then transferred into H_2_O *via* a glass Pasteur pipette and gently mixed by pipetting up and down. The vacuum pump (−3 mbar) was then started and the wells allowed to dry. The wells were then washed twice with 15% ACN in ultrapure H_2_O (500 µL, 0.1% v/v formic acid). The pressure was then reduced to −15 mbar and the wells dried for 15 minutes. The waste container under the 96-well sorbet plate was then replaced with 750 µL silylated glass insert vials in a deep 96-well plate. The endocannabinoids were eluted by allowing acetone (350 µL, 0.1% v/v formic acid) to pass through the sorbent bed twice. The samples were then evaporated to dryness under a stream of N_2_ prior to resuspension in a 70% solution of ACN (100 µL, 30% ultrapure water, 1% v/v acetic acid). The samples were then vortexed briefly and stored at −20 °C prior to the LC-QqQMS experiment.

The endocannabinoids were separated using ultra-high-performance liquid chromatography (UHPLC). The two eluent solvents were (i) H_2_O (1% v/v acetic acid) and (ii) ACN. The gradient was set up as follows: the initial conditions were 70% B for 4.5 minutes increasing to 80% B by 7 minutes. By 8 minutes, the gradient was increased to 95% B and held there for 10 minutes. The gradient was reduced to the starting conditions by 10.5 minutes and the column flushed with a minimum of 10 column volumes. The flow rate was set at 0.5 mL/min throughout the run and the column temperature set to 60 °C. The column used for the separation was a Waters ACQUITY BEH reverse phase C18 column (130 Å, 1.7 µm, 2.1 mm × 50 mm). The injection volume was 10 µL. The needle was washed three times both before and after the injection with two solvents: (i) IPA:ACN (1:1), and (ii) H_2_O:MeOH (1:1). The mass spectrometer 5500 QTRAP (AB Sciex Inc., Framingham, MA) was set up in scheduled multiple reaction monitoring (MRM) mode which allows for the setup of detection windows around the peak of interest to improve sensitivity. The details of the MRM transitions can be found in Supplementary Table 1.

### Statistical Analysis

The non-parametric Mann-Whitney U test was used for comparing the levels of circulating endocannabinoids, and performed using GraphPad Prism v. 7.04 (GraphPad Software Inc., San Diego, CA). In order to compare the association between circulating endocannabinoids with brain CB1R availability, two methods were utilized: (i) partial least squares regression (PLS-R) modeling was used in order to combine multiple brain regions and circulating endocannabinoid measures. This analysis was performed using PLS-Toolbox v. 8.6 (Eigenvector Research Inc., Manson, WA) and MATLAB 2017b (Mathworks Inc., Natick, MA). Due to the small number of samples, a leave-one-out cross-validation, using the number of iterations of the smallest group size, was utilized to ensure model validity. (ii) Spearman correlation coefficients were calculated using the statistical toolbox in MATLAB 2017b and p-values < 0.05 (two-tailed) were considered significant for the correlations. The individual spearman correlation coefficients (R) were illustrated as a heatmap using the ‘corrplot’ package for the R statistical programming language^51^. The individual correlations between the serum endocannabinoids and CB1R availability were plotted and analyzed using GraphPad Prism.

### Generation of the statistical association brain maps

In order to visualize the associations of CB1R availability across different brain regions, the R^2^ values from the individual brain regions arising from the PLS-R modelling were mapped onto the same brain template as the PET image analysis using MATLAB 2017b. The resultant image was then visualized using CARIMAS v. 2.9 (Turku PET Centre, Turku, Finland). The color scale for each brain region was defined by the healthy controls in each cohort and then applied to the FEP patients.

## Results

### Levels of circulating endocannabinoids

We measured a total of ten endocannabinoids and related structures (Supplementary Table 1) from serum of both HC and FEP patients from the two study sites. Considerable increases in AG were observed in some samples, which can be explained by variable freezing times^52^. These samples were excluded from subsequent analyses, resulting in the following available sample numbers: Turku (n = 10, HC; n = 8, FEP) and London (n = 11, HC; n = 15, FEP). In the Turku study, there was a reduction in circulating levels of arachidonic acid (AA; 140.6 ± 29.9 vs. 104.1 ± 26.8 ng/mL) and OEA (3.23 ± 1.04 vs. 2.29 ± 0.50 ng/mL) in the FEP patients (Figure 2a). AA forms the backbone of two major endocannabinoids (AEA and 2-AG)^53^, and is released when they are broken down by fatty acid amide hydrolase (FAAH)^54^ and monoacylglycerol lipase (MGL)^55^. Therefore, the decreased levels of AA in FEP suggest reduced breakdown of these two metabolites. In line with this, there is a trend towards an increase in AG observed in the FEP group (2.32 ± 0.94 vs. 3.69 ± 2.02 ng/mL). Due to the use of serum, it was, however, not possible to differentiate between the levels of 1-AG and 2-AG, because of their rapid isomerization at room temperature. Therefore, only the concentrations of total AG are reported.

**Figure 1.**
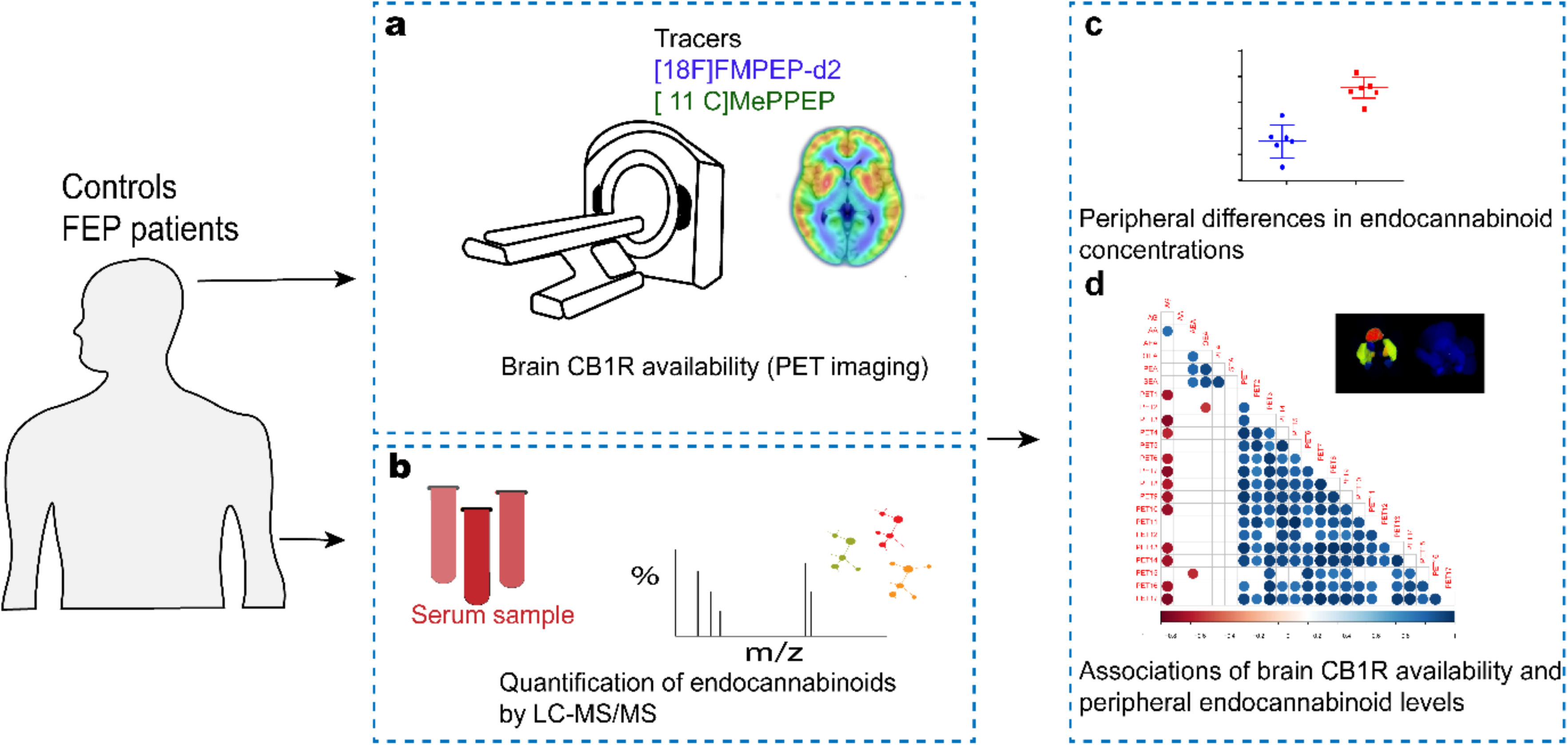
An overview of the experimental design to study associations between brain cannabinoid receptor 1 (CB1R) availability and circulating endocannabinoids. (**a**) CB1R availability was investigated *in vivo* in male patients with, first episode psychosis (FEP) and healthy controls (HC) using a CB1R-selective radiotracer using positron emission tomography (PET) with arterial blood sampling. This study was performed at two study sites using two independent samples in the city of Turku, Finland and an inner-city area of London, United Kingdom. A [^18^F]FMPEP-d2 PET tracer was used in Turku while [^11^C]MePPEP tracer was used in London. (**b**) Quantification of circulating endocannabinoids was performed in matched FEP and HC subjects using a quantitative liquid-chromatography triple-quadrupole mass spectrometry assay. (**c**) Peripheral differences in endocannabinoids between FEP and HCs were evaluated by univariate statistics. In order to assess if circulatory endocannabinoids associated with CB1R availability in the brain, we performed statistical correlation analysis between six endogenous endocannabinoids and seventeen CB1R tracer distribution volumes. The correlation analysis was performed separately between FEP and HC for the two study sites.

**Figure 2.**
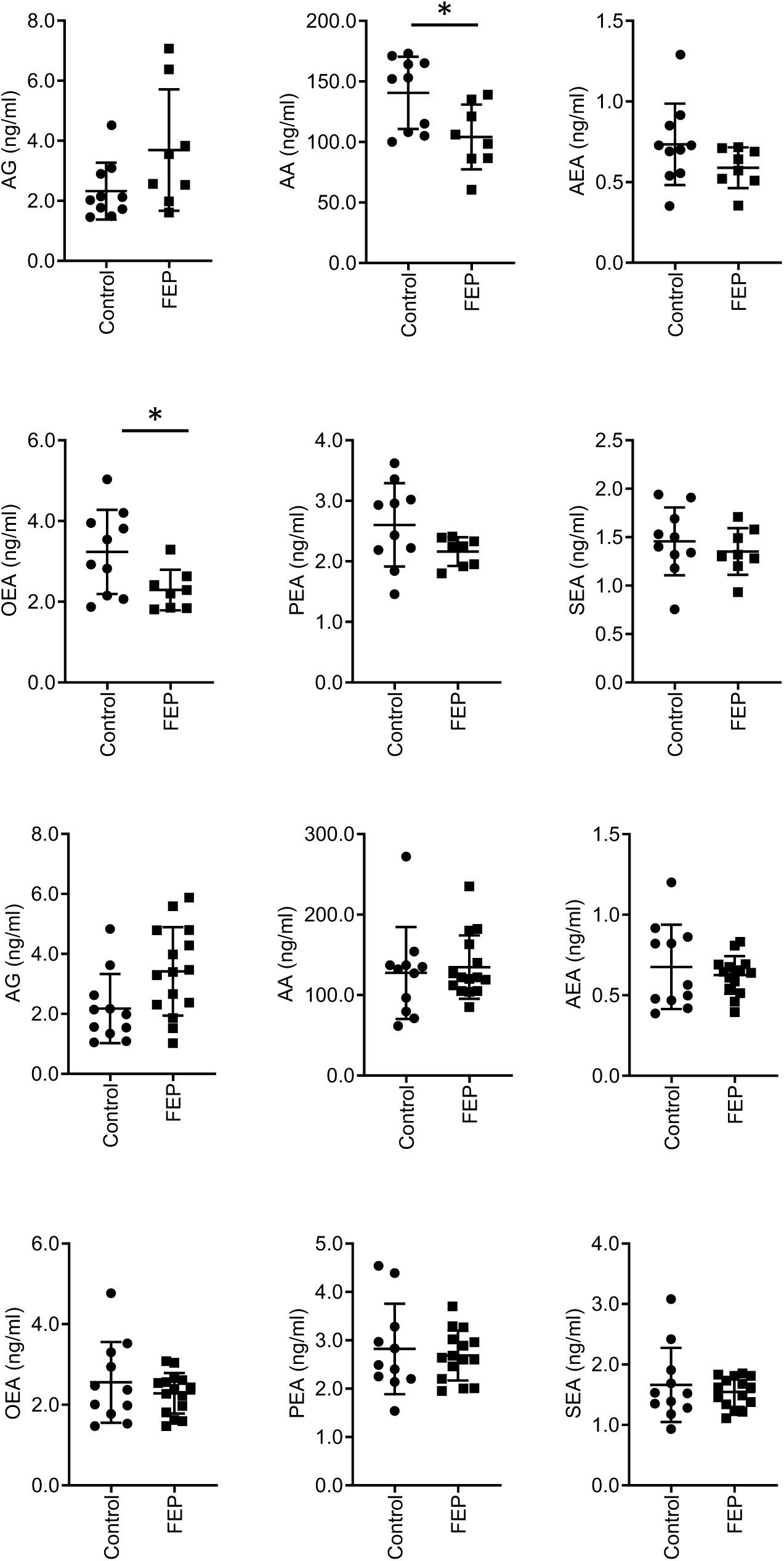
Scatter plots of the levels of circulating endocannabinoids and related structures from both the Turku (top six) and KCL (bottom six) cohorts. *p < 0.05.

The same endocannabinoids were, however, not altered in patients with FEP in the London study (Figure 2b). In Turku, patients had already started antipsychotic medication, whilst in London, the patients were mostly drug-naïve or free of pharmacological treatments^27^. Previous literature has shown that endogenous central elevations in anandamide (a CB1R agonist) found in patients with FEP^14, 56^ are absent in patients taking antipsychotics^56^. Thus, antipsychotic treatment in the Turku sample may account for the differences we observe. However, this discrepancy may also be due to other differences between the study populations, such as diagnosis, substance use, diet or duration of illness.

### Associations between central and peripheral endocannabinoid systems

Given the known role of the endocannabinoid system in dietary regulation^57^ and weight gain^58^, we next investigated the associations between peripheral endocannabinoid levels and CB1R availability in the ROIs described in the methods.

Initially, we explored how circulating levels of endocannabinoids are associated with CB1R availability in the brain in both studies using PLS regression, which allowed us to see how all of the endocannabinoids are associated with individual brain regions in one model. The R^2^ values were increased in healthy controls both in the Turku (Figure 3a) and London (Figure 3c) cohorts, as compared to the FEP group (Figure 3b, Turku and Figure 3d, London). We next explored the univariate associations between levels of circulating endocannabinoids and CB1R availability in the brain. Positive correlations of CB1R availability between the brain regions were observed in HCs from both study sites (Figure 3e). Similarly, levels of circulating endocannabinoids were also positively correlated with each other in the healthy subjects (Figure 3e). Interestingly, the levels of peripheral endocannabinoids and central CB1R availability were, conversely, negatively correlated in HCs in both Turku and London samples (Figure 3e). In FEP subjects, CB1R availability was also positively correlated between the different brain regions (Figure 3f). However, the degree of positive correlation between the circulating endocannabinoids was reduced (Figure 3f).

**Figure 3.**
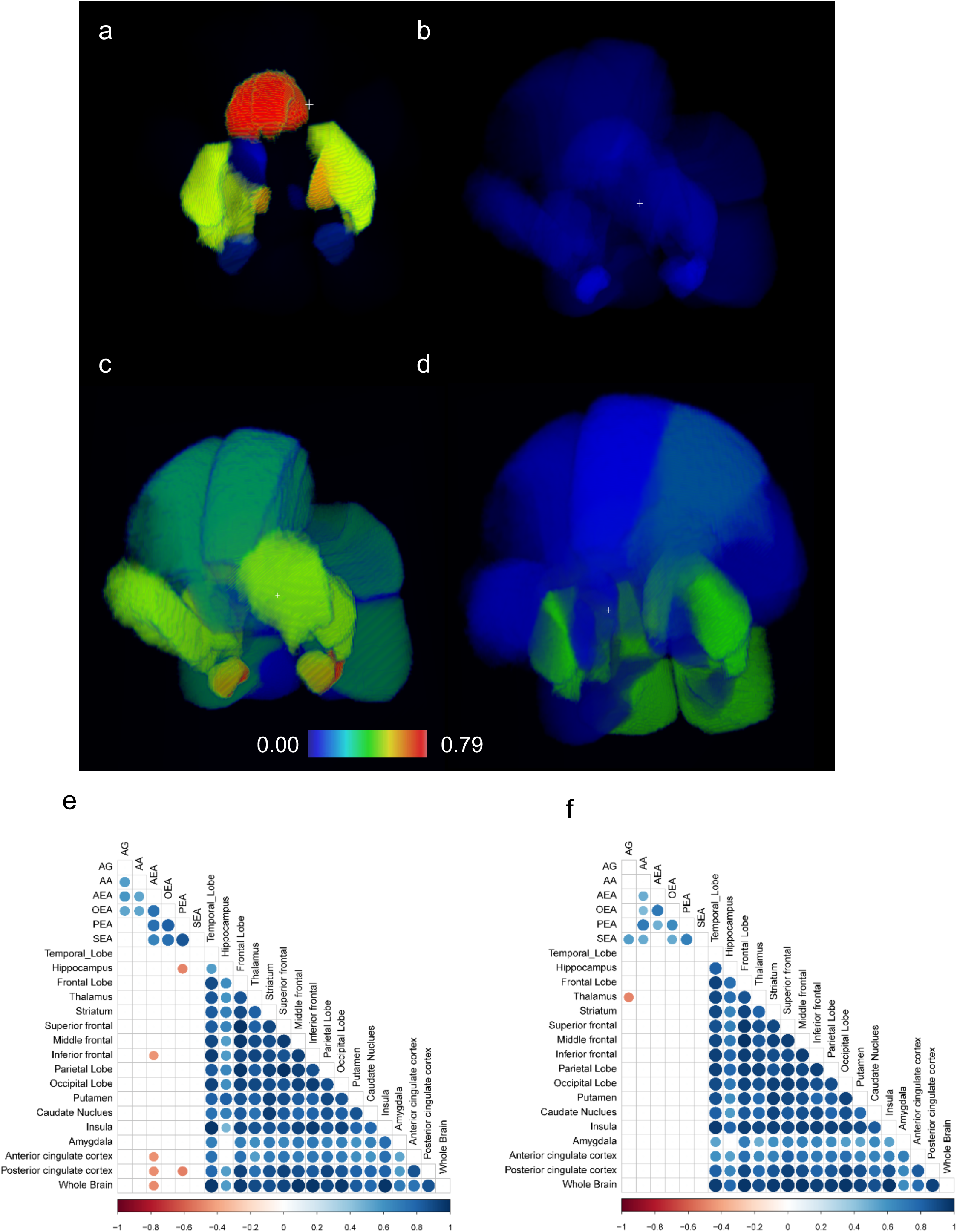
Correlation analysis, performed between peripheral endocannabinoid concentrations and the distribution volumes of brain CB1R availability separately for patients with FEP and HC. Correlation brain maps where the R^2^ values from the PLS regression analysis are projected into the 3D brain regions used for the PET analysis in the healthy controls (**a**, Turku, **c**, London) and FEP patients (**b**, Turku, **d**, London). Red shading denotes high R^2^, blue low R^2^. (**e**) Spearman correlation coefficients between CB1R availability and circulating endocannabinoids in HC from both Turku and London. (**f**) Spearman correlation coefficients between CB1R availability and circulating endocannabinoids are shown in HC and (**f**) FEP from both Turku and KCL. The CB1R availability was combined by first autoscaling the data from each site. The correlation coefficient values revealed positive correlations between CB1R availability in different regions of the brain in both FEP and HC. There was significant negative association between peripheral AEA and PEA concentrations and CB1R availability in certain regions of the brain among HCs, and this relationship was absent in FEP patients, at each respective site. Here, the blue gradient represents positive correlation while the red gradient indicates negative correlation (p-values < 0.05).

Notably, the negative correlations between CB1R availability and levels of circulating endocannabinoids, as observed in HC, were absent in FEP patients (Figures 3e and 3f). The loss of this association is more pronounced in the Turku study (Supplementary Figure 1a-b) compared to the London study (Supplementary Figure 1c-d). This relative lack of association between central CB1R availability and circulating endocannabinoid levels in FEP suggests that there may normally be a functional link between these two systems, which may be dysregulated in FEP.

In order to further examine the associations between central CB1R availability and circulating peripheral endocannabinoids, we selected the brain region, from the available ROIs, with the highest R^2^ value from the PLS-R modeling; the posterior cingulate cortex (Figure 3). For the majority of serum endocannabinoids measured in FEP patients, there was a clear loss of any negative correlation with CB1R availability, with few positive associations observed in the FEP patients (Figure 4). This effect was more pronounced in the Turku study (Supplementary Figure 2a), with similar trends observed in the London study, though these did not reach statistical significance (Supplementary Figure 2b).

**Figure 4.**
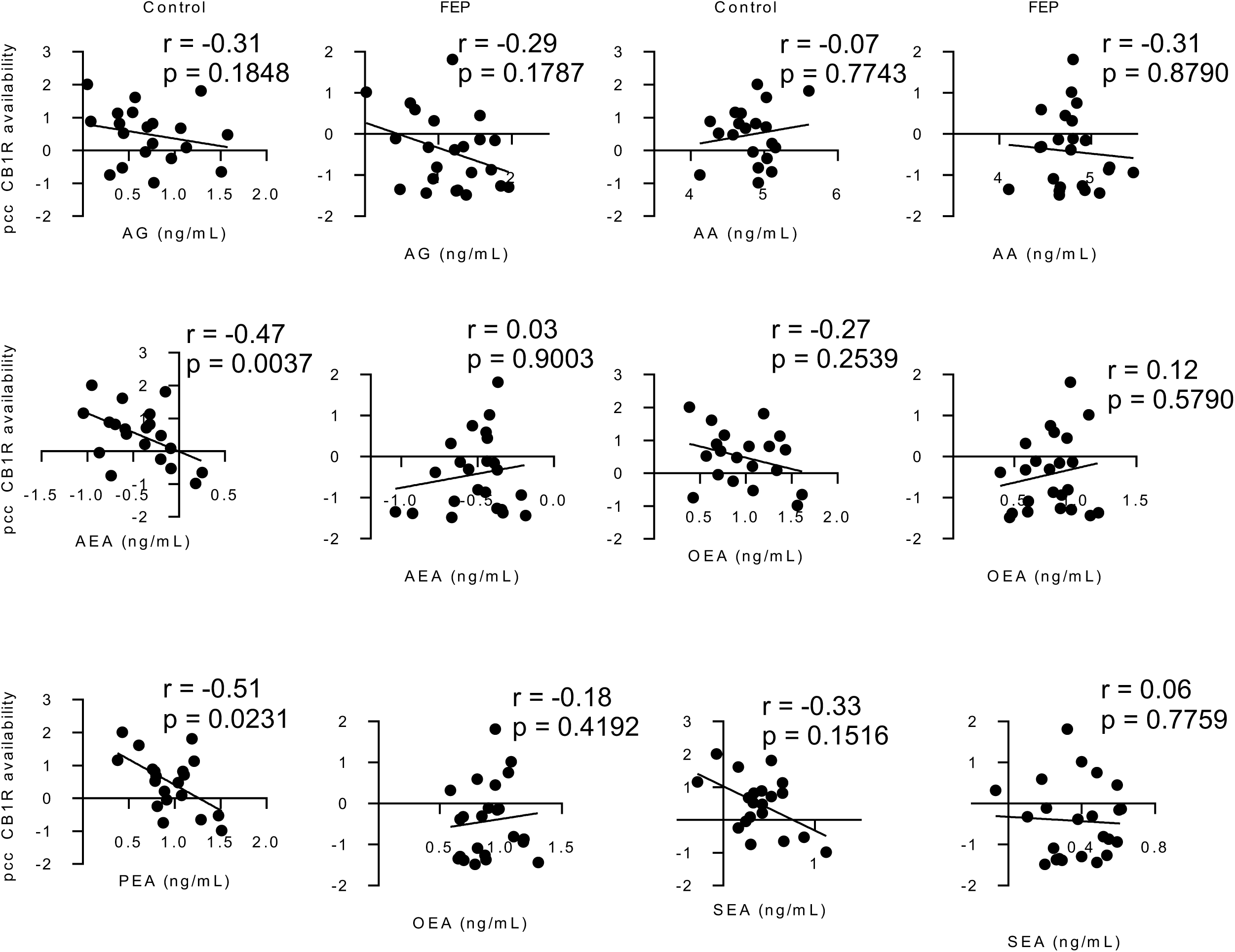
Scatter plots fitted with a linear regression model of CB1R availability (combined data from both studies, scaled to zero mean and unit variance within each study separately) in the posterior cingulate cortex (PCC), versus the log-transformed circulating levels of endocannabinoids. The line shows the linear model for each dataset.

## Discussion

We demonstrated that the cross-talk between the endocannabinoid networks in the brain and periphery are dysregulated in FEP, as compared to healthy controls. Reduced levels of AA and OEA in FEP subjects were observed in the Turku study, but not in London, where the patients were predominately drug-naïve or free from pharmacological treatments. A negative association between brain CB1R availability in the majority of the ROIs used in the PET imagiing and serum endocannabinoid levels was observed in healthy individuals, but not in FEP patients across the two independent cohorts of patients.

Endocannabinoids are lipid mediators with a vital role in the maintenance of metabolic and immune homeostasis^59, 60^. Several studies have found that systemic lipid profiles are dysregulated in various psychotic disorders^9, 61^. Our study corroborates these earlier findings, suggesting that the systemic ECS is dysregulated in psychosis^12, 29, 62^. In the periphery, a significant drop in the circulatory endocannabinoid, anandamide, was previously found to accompany the acute phase of schizophrenia^12^. This finding is at odds with findings from the London study as well as previous literature showing that patients with FEP not taking antipsychotics or using cannabis show increased levels of anandamide in cerebrospinal fluid^14, 29, 56^. However, the finding from the Turku study is consistent with previous work showing a downregulation of enzymes involved in the synthesis of endocannabinoid ligands in peripheral blood mononuclear cells (PBMCs) of FEP patients, as compared to matched, healthy controls^62^. In addition to FEP, other neuroinflammatory diseases including multiple sclerosis, Huntington’s and Parkinson’s diseases, are characterized by ECS alterations in the periphery^63^.

Our previous data links rapid weight gain in FEP with elevated baseline levels of triglycerides of low carbon number and double bond count ^9^, which are known to be associated with *de novo* lipogenesis, inhibition of lipolysis, and NAFLD^64, 65^. Endocannabinoids are known to stimulate *de novo* lipogenesis and obesity^66^. In our previous hepatic venous catheterization study in humans, we found elevated liver fat to be associated both with increased levels of 2-AG as well as higher splanchnic production of triglycerides of low carbon number and double bond count, suggesting that the ECS could contribute to the development of human NAFLD by affecting hepatic lipid metabolism^64^.

OEA is the monounsaturated analogue of AEA, but, unlike AEA, acts independently of the cannabinoid pathway, with no reported direct activity at the two known cannabinoid receptors, CB1R and CB2R^67^. OEA stimulates fat utilization as a peroxisome proliferator-activated receptor alpha (PPARα) agonist^68^. Activation of PPARα by OEA is known to induce satiety and regulate body weight^69^. A study in mice also suggests that OEA inhibits food intake by recruitment of the brain histaminergic system^70^. Our data suggest that decreased OEA in FEP subjects is likely the effect of antipsychotic meditation, as this was not seen in the cohort that was predominately unmedicated. Inhibition of lipolysis due to diminished OEA^69^ may also contribute to increased liver fat, *i.e.*, the accumulation of triacylglycerols in the liver, as observed by lipidomics in non-obese FEP patients who subsequently rapidly gained weight^9^. It is thus plausible that OEA levels are an early marker of propensity for FEP patients to gain weight. This hypothesis will clearly need to be examined prospectively, in a larger study setting.

Our study also shows that peripheral endocannabinoids and central CB1R availability are inversely correlated in healthy individuals, while no such association is observed in FEP patients. The CB1R modulates energy homeostasis *via* both central and peripheral mechanisms^71^. The association between peripheral and central ECS measures suggests common systemic regulation of ECS in healthy individuals, which, however, is absent in FEP. Our findings from the Turku cohort indicate that both central and peripheral measures of the endocannabinoid system were altered in patients with affective / non-affective psychosis who were taking antipsychotic medication. In contrast, our findings from the London cohort indicate that patients who were predominately medication naïve/free from pharmacological treatments, show central alterations in CB1R availability without any alterations in peripheral levels of endocannabinoids. This suggests that peripheral measures of endocannabinoids may not be a useful biomarker for indexing central endocannabinoid dysfunction in schizophrenia, although, alternatively, this negative finding may also be due to the small sample size, or differences in sample clinical characteristics between studies including diagnosis, illness severity and illness duration.

Taken together, our data provides evidence of ECS dysregulation in FEP both centrally^27^ and in the periphery, in medicated patients with FEP. Our findings should be considered as preliminary due to our small sample size and partial inconsistency of results. Further studies are clearly needed in order to further examine the cross-talk between the peripheral and central ECSs in health and psychosis.

## Acknowledgements

This project has received funding from the European Union’s Seventh Framework Programme for project METSY—Neuroimaging platform for characterization of metabolic co-morbidities in psychotic disorders (no. 602478). The authors thank to Aidan McGlinchey for assistance with editing the manuscript.

## Author contributions

M.O., J.H., and O.H. initiated, designed, and supervised the study. A.D. T.R., T.L., and T.H. acquired serum endocannabinoid data by mass spectrometry. B.F., H.L., T.M., and M.V. acquired neuroimage data. A.D., B.F., H.L., and M.O. analyzed the data. A.D., S.L. and M.O. wrote the first draft of the manuscript. All authors approved the final version.

---

**Supplementary Table 1.** Tandem MS analysis of endocannabinoids. Information on quantification and selective reaction monitoring (SRM) transitions for identification.

| Compound | Ionization | Ion Type | Retention Time | Internal Standard | SRM (amu) | CE (eV) |
| --- | --- | --- | --- | --- | --- | --- |
| THC-COOH | Negative | Quantifier | 1.23 | THC-COOH d9 | 343.400 → 245.100 | -32 |
| THC-COOH | Negative | Qualifier | 1.23 | THC-COOH d9 | 343.400 → 299.000 | -32 |
| PEA | Positive | Quantifier | 3.13 | AEA-d8 | 300.400 → 62.100 | 19 |
| PEA | Positive | Qualifier | 3.13 | AEA-d8 | 300.400 → 283.100 | 19 |
| 1-AG | Positive | Quantifier | 3.24 | 2-AG d5 | 379.300 → 287.300 | 21 |
| 1-AG | Positive | Qualifier | 3.24 | 2-AG d5 | 379.300 → 203.200 | 21 |
| 2-AG | Positive | Quantifier | 2.97 | 2-AG d5 | 379.300 → 287.300 | 21 |
| 2-AG | Positive | Qualifier | 2.97 | 2-AG d5 | 379.300 → 203.200 | 21 |
| 2-AGe* | Positive | Quantifier | 3.23 | 2-AG d5 | 365.400 → 273.200 | 20 |
| 2-AGe* | Positive | Qualifier | 3.23 | 2-AG d5 | 365.400 → 121.000 | 20 |
| NADA* | Positive | Quantifier | 2.69 | NADA d8 | 440.300 → 137.100 | 30 |
| NADA* | Positive | Qualifier | 2.69 | NADA d8 | 440.300 → 154.100 | 30 |
| AEA | Positive | Quantifier | 2.26 | AEA-d8 | 348.300 → 62.100 | 32 |
| AEA | Positive | Qualifier | 2.26 | AEA-d8 | 348.300 → 133.100 | 32 |
| OEA | Positive | Quantifier | 3.61 | AEA-d8 | 326.300 → 62.100 | 20 |
| OEA | Positive | Qualifier | 3.61 | AEA-d8 | 326.300 → 309.300 | 20 |
| AA | Negative | Quantifier | 4.48 | AA-d8 | 303.300 → 259.100 | -20 |
| AA | Negative | Qualifier | 4.48 | AA-d8 | 303.300 → 58.900 | -20 |
| SEA | Positive | Quantifier | 5.92 | AEA-d8 | 328.400 → 62.100 | 23 |
| SEA | Positive | Qualifier | 5.92 | AEA-d8 | 328.400 → 311.400 | 23 |

\*Analyte below limit of detection in study samples.

**Supplementary Figure 1.**
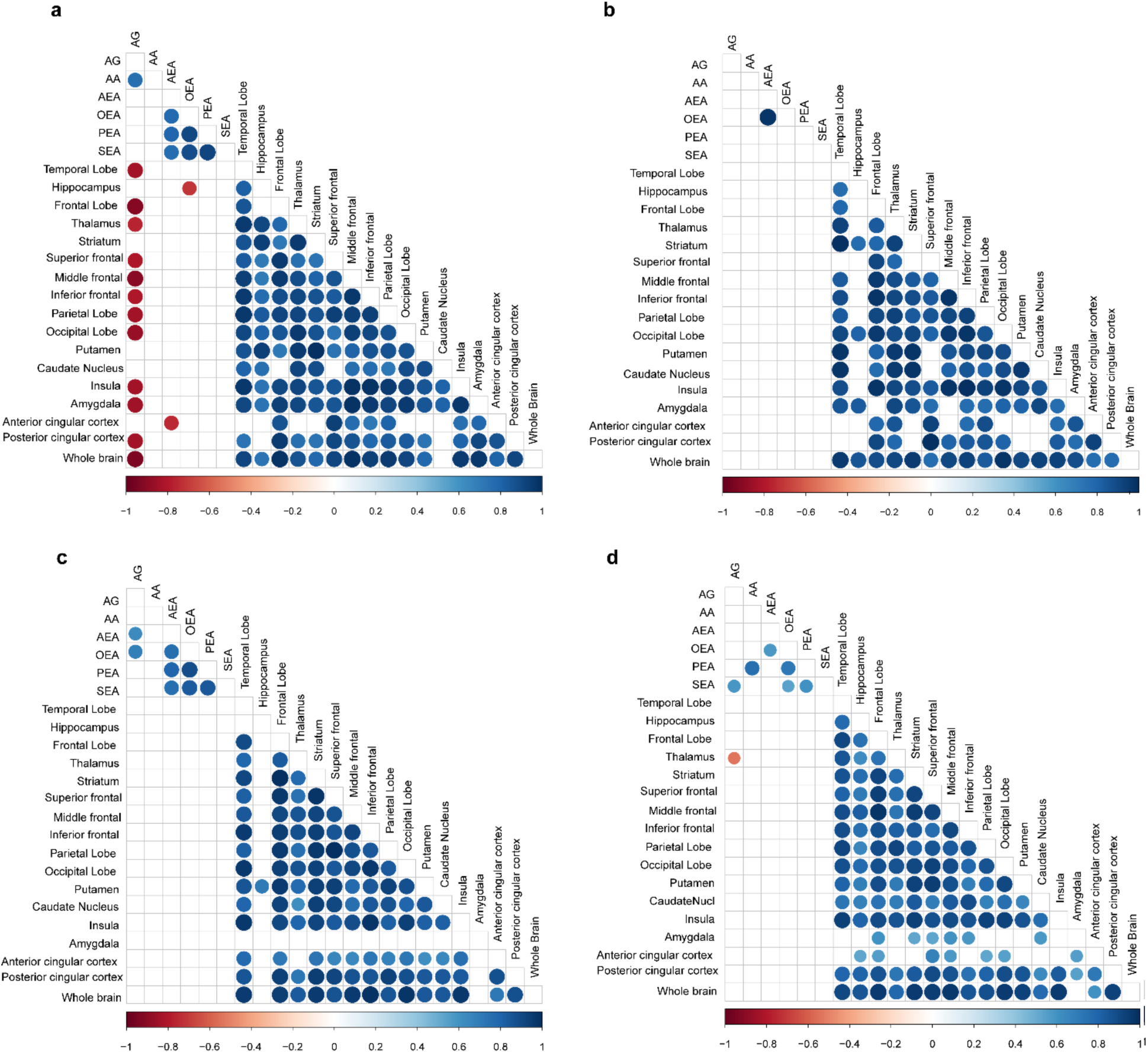
Correlation analysis, performed between peripheral endocannabinoid concentrations and the distribution volumes of brain cannabinoid receptor 1 availability separately between FEP patients HC at the two study sites (**a**) Spearman correlation coefficients between CB1R availability and circulating endocannabinoids in HC from the city of Turku. (**b**) Spearman correlation coefficients between CB1R availability and circulating endocannabinoids in FEP from the city of Turku. The correlation coefficient values revealed positive correlation between CB1R availability in different regions of the brain in both FEP and HC. (**c**) Spearman correlation coefficients between CB1R availability and circulating endocannabinoids in HC from an inner-city area of London. (**d**) Spearman correlation coefficients between CB1R availability and circulating endocannabinoids in FEP from inner city area of London. There was significant negative association between peripheral arachidonoyl glycerol (1+2) concentrations and CB1R availability in certain regions of the brain among HCs, which was absent in FEP patients, at different sites, respectively. Here, the blue gradient represents positive correlation while the red gradient indicates negative correlation (p-values < 0.05).

**Supplementary Figure 2.**
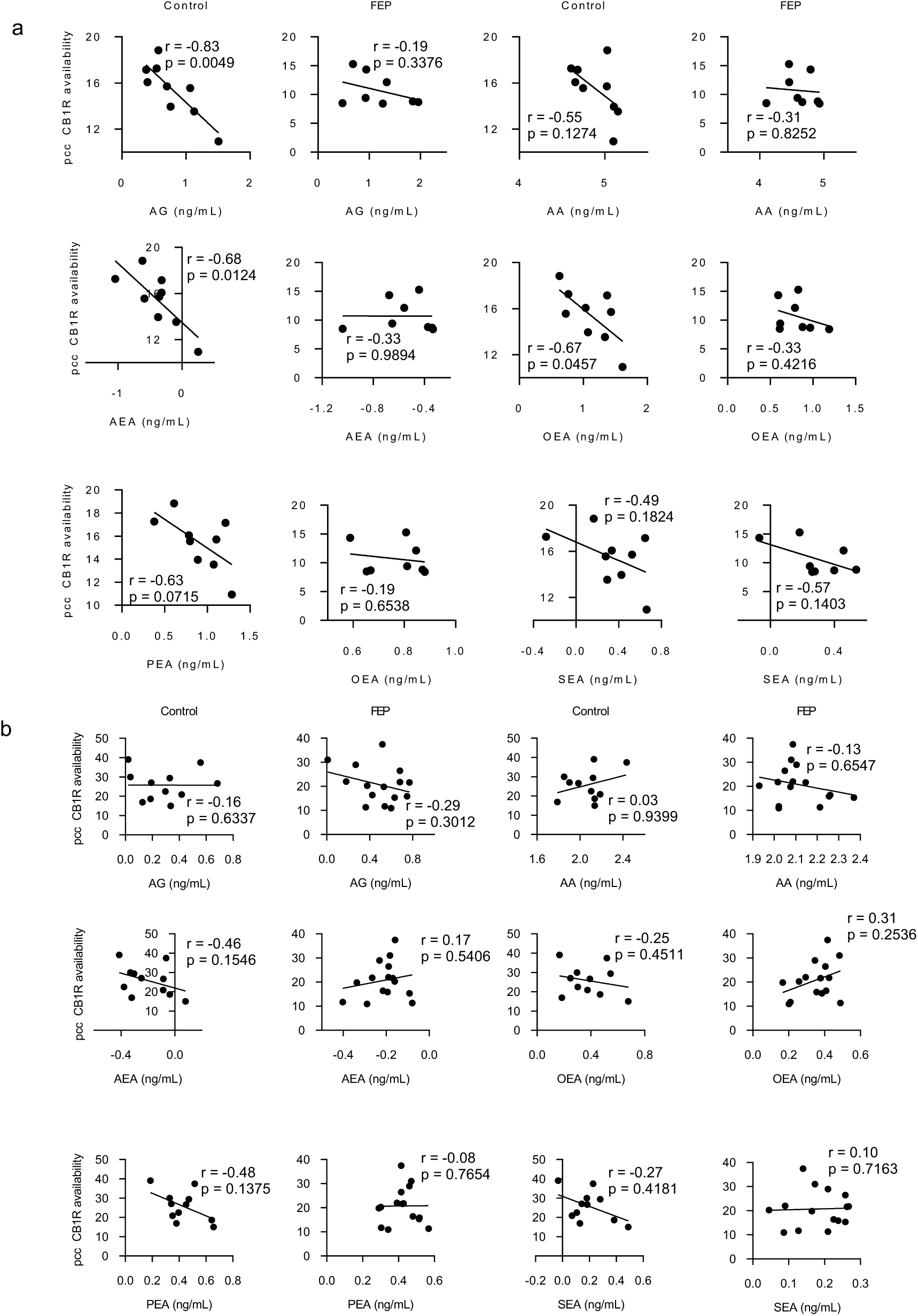
(**a**) Scatter plots fitted with a linear regression model of CB1R availability in the posterior cingulate cortex (PCC) versus the log-transformed circulating levels of endocannabinoids from the Turku cohort. (**b**) Scatter plots fitted with a linear regression model of CB1R availability in the posterior cingulate cortex (PCC) versus the log-transformed circulating levels of endocannabinoids from the London cohort. The line shows the linear model for each dataset.

